# Clove Aqueous Extract Triggers a Multi-Organellar Stress Crisis through Lysosomal Destabilisation and Mitochondrial Hyperpolarisation to Suppress Patient-Derived Ovarian Cancer Cells

**DOI:** 10.64898/2026.01.28.702206

**Authors:** Yazid Ghanem, Hadeel Odwan, Musha Yang, Victoria Malone, Fawza Alenazi, Fears Abu Saadeh, Steven G. Gray, Derek Doherty, Cara Martin, Sharon O’Toole, John J. O’Leary, Bashir M. Mohamed

## Abstract

Ovarian cancer (OC) remains a lethal malignancy with limited therapeutic options, underscoring the need for the identification of novel agents. Natural products like clove (*Syzygium aromaticum*) have shown promising anti-cancer activity, but their mechanism in OC is poorly understood. This study investigates the anti-tumour effects and underlying mechanisms of a clove aqueous extract (CAE) on a panel of patient-derived OC cells. We found that CAE significantly inhibited cellular proliferation and induced cell death in a time-and dose-dependent manner. Mechanistically, CAE induced profound cellular stress, activating the transcription factor ATF-2. This was accompanied by a significantly increased lysosomal stress response, as evidenced by increased lysosomal mass/acidity, and a pathogenic hyperpolarisation of the mitochondrial membrane potential (ΔΨm). The bioenergetic crisis induced as a consequence resulted in a sharp reduction in cellular oxygen consumption rate (OCR). Notably, the sensitivity to CAE-induced lysosomal and mitochondrial dysfunction varied across cell lines, revealing distinct phenotypic responses. Our results demonstrate that clove extract exerts its anti-tumour effects by orchestrating a multi-organellar stress response, positioning lysosomal disruption as a central event in its mechanism of action. This study provides a strong rationale for the further development of clove-based interventions for OC.

## Introduction

Ovarian cancer (OC) remains the most lethal gynaecologic malignancy worldwide, and despite advances in treatment, survival outcomes have improved only modestly. In the United States, recent surveillance estimates project approximately 20,890 new cases and 12,730 deaths in 2025 (1). The 5-year relative survival rate remains at roughly 50–55%, reflecting only incremental gains over several decades (1–3). A major contributor to the persistently poor prognosis is the fact that most patients are diagnosed at an advanced stage; nearly 70% present with FIGO stage III or IV disease. This is largely due to the nonspecific nature of early symptoms and the continued absence of an effective screening strategy for the general population (4–7).

The current standard first-line treatment for OC consists of maximal cytoreductive (debulking) surgery followed by platinum-based chemotherapy, most commonly carboplatin or cisplatin in combination with paclitaxel (8–11). Although initial response rates can approach 80%, more than half of patients eventually experience disease recurrence, primarily due to the development of intrinsic or acquired resistance to chemotherapy (8–11). Moreover, platinum and taxane regimens are also associated with cumulative toxicities. Nephrotoxicity and peripheral neurotoxicity can become dose-limiting and may restrict further treatment options (12–15). Collectively, these challenges highlight the urgent need for novel therapeutic strategies that act through alternative mechanisms and exhibit improved safety profiles. Natural products have historically played a central role in cancer drug discovery, with an estimated 60% of currently used chemotherapeutic agents derived from natural compounds or their structural analogues (16–18). Paclitaxel, a cornerstone of OC therapy, is itself derived from the Pacific yew tree, underscoring the enduring relevance of plant-based molecules in oncology (19,20). Beyond their direct cytotoxic effects, natural products are increasingly being investigated for their ability to overcome chemoresistance and mitigate treatmentrelated toxicities (16,21).

Clove (Syzygium aromaticum L.), a spice widely used in traditional Chinese medicine, Ayurvedic medicine, and Kampo practices, contains a diverse array of bioactive constituents, including eugenol, β-caryophyllene, flavonoids, and various triterpenoids (22–24). Clove extracts have demonstrated antioxidant, anti-inflammatory, antimicrobial, and anticancer activities in various experimental models (25–30). Of particular relevance to OC, the clove derived flavonoid kumatakenin has been shown to induce apoptosis and inhibit tumour associated macrophage activation in preclinical studies (31,32). More broadly, clove extracts exert cytotoxic and pro-apoptotic effects across multiple cancer types, largely through modulation of cell-cycle regulation, DNA damage responses, and mitochondrial stress pathways (25–32). Given the metabolic vulnerabilities and organelle-specific dysfunctions characteristic of OC cells, we hypothesised that clove aqueous extract (CAE) may exert antitumor effects by coordinating stress responses within both lysosomes and mitochondria. To test this hypothesis, we employed a panel of patient-derived OC models representing major epithelial subtypes, including high-grade serous ovarian cancer (HGSOC) and carcinosarcoma, thereby capturing the molecular and clinical heterogeneity of the disease (33–36).

This experimental approach allowed us to determine whether CAE exhibits broad antitumor activity across OC subtypes or displays selective efficacy, to delineate the temporal sequence of lysosomal and mitochondrial disruption, and to identify the downstream pathways leading to cell death. Together, these findings provide mechanistic insight into the anticancer activity of clove-derived compounds and support the potential of organelle-targeting strategies as a novel avenue for improving OC treatment.

## Materials and methods

### Ethical Approval

This study was approved by St. James’s Hospital and Adelaide and Meath Hospital, Dublin, incorporating the National Children’s Hospital Research Ethics Committee (Reference: 2012/11/04). All procedures were conducted in accordance with the Declaration of Helsinki and relevant institutional and national guidelines. Written informed consent was obtained from all participants or their legal guardians prior to sample collection.

### Patient-Derived OC Cells Isolation and Expansion

Patient-Derived OC cells with high-grade serous (HGSOC) and carcinosarcoma subtypes were isolated from either ascites fluid obtained from three patients (designated OCAS12, OCAS14 and OCAS17) and/or from ovarian tumour tissue (OCAST16) and used throughout this study. As previously described (37), cells were maintained in RPMI 1640 medium (GIBCO, Invitrogen, Ireland) supplemented with 10% (v/v) fetal calf serum, 20 mM HEPES, 10 μM nicotinamide, 10 μM SB202190, 1.25 mM N-acetyl-L-cysteine, 10 ng/mL FGF-10, 1 ng/mL FGF-2, 1× B27 supplement, Primocin (1:100, v/v), 10 μM Y-27632, 2 mM Lglutamine, and 100 U/mL penicillin-streptomycin. Cultures were maintained at 37°C in a humidified incubator with 5% CO_2_. Four days after initial plating, non-adherent cells were removed by washing with phosphate-buffered saline (PBS), and fresh culture medium was added. Cells were subsequently expanded for experiments under standard conditions.

### Patient-Derived OC Cell Phenotypic Characterisation

Patient-Derived OC cells were characterised for cancer-associated and epithelial–mesenchymal transition (EMT) markers, including mucin-1 β-catenin, ErbB-3, EGFR, claudin7, folate receptor-α, and mesothelin. Cells were seeded at 1 × 10 cells per well in 96well plates and incubated overnight at 37°C with 5% CO_2_. The following day, cells were washed with PBS and fixed in 3% paraformaldehyde (PFA; Sigma-Aldrich, Ireland) for 20 minutes at room temperature (RT). After fixation, cells were blocked with 5% bovine serum albumin (BSA; Sigma-Aldrich, Ireland) for 1 hour at RT.

Cells were then incubated overnight at 4°C with mouse monoclonal primary antibodies (1:200; Santa Cruz Biotechnology, Germany). The next day, cells were washed with PBS and incubated with a goat anti-mouse FITC-conjugated secondary antibody (1:300; ThermoFisher Scientific, Dublin, Ireland) for 1 hour at RT. Nuclear staining was performed using Hoechst 33342 (1:1000; ThermoFisher Scientific, Dublin, Ireland) for 20 minutes at RT. Following final PBS washes, cells were imaged using an EVOS inverted fluorescence microscope (Figure 1).

**Figure 1:**
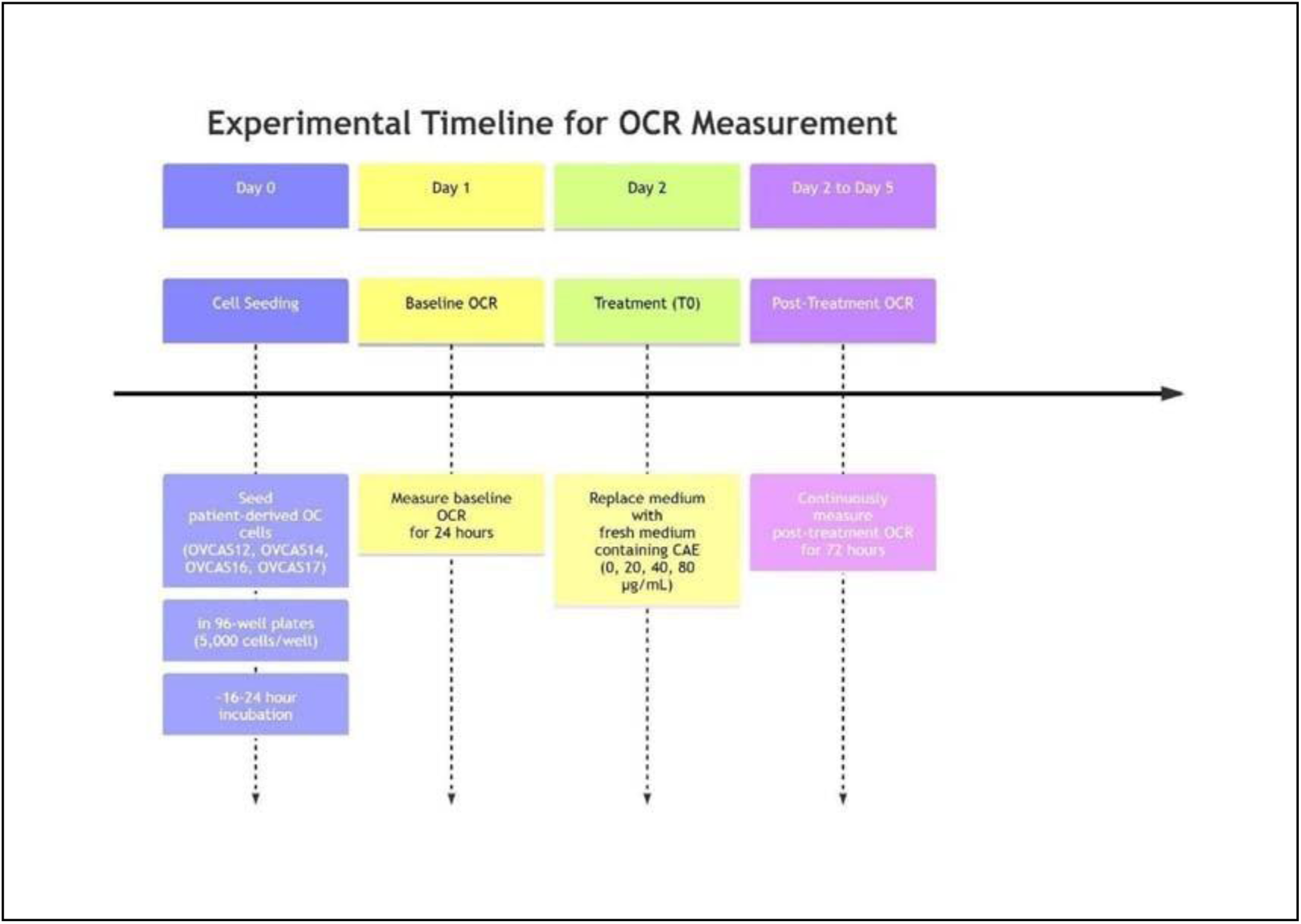
Experimental Workflow for Real-Time OCR Profiling: This diagram illustrates the sequential steps and timeline for measuring the Oxygen Consumption Rate (OCR) in response to clove extract (CAE) treatment.

### Construction and treatment of scaffold-free PDOs

All Patient-Derived OC Cells (OCAS12,OCAS14 OCAS17 and OCAST16 cells) were seeded in a and fed with DMEM/RPMI (Invitrogen, Ireland) supplemented with 10% (v/v) foetal calf serum, 20 mM HEPES, 10 µM SB202190, 1.25 mM N-Acetyl-L-cysteine, 10 ng/mL FGF-10, 1 ng/mL FGF-2, 1X B27 Additive, 1:100 (v/v) Primocin, and 10 µM Y27632, 2 mM Lglutamine and 100 U/mL penicillin-streptomycin. The cells were maintained at 37°C in humidified air with 5% CO_2_ until spherical organoids formed within approximately eight days.

## Preparation of Clove Crude Aqueous Extract (**CAE)**

Dried clove buds (*Syzygium aromaticum L.*) were purchased from a local commercial supplier (Dublin). The buds were finely ground using a Krups coffee grinder. For each extraction, 5 g of ground clove powder was suspended in 50 mL of cell culture medium (RPMI). The mixtures were incubated on a Stuart Scientific digital roller mixer for 24 hours at room temperature. Following incubation, samples were centrifuged at 5,000 × g for 30 minutes at 4°C using a Hettich Rotina 35 R refrigerated centrifuge. The resulting supernatants of CAE were collected and filtered under sterile conditions through a 0.22 µm PES membrane filter (Millex-GP, Merck) using a 50 mL Terumo syringe. Filtered CAE were transferred into sterile 50 mL Sarstedt screw-cap tubes and stored at 4°C until use.

### Cytotoxic effects of CAE

#### Assessment of Cell Viability using CCK-8 Assay

To measure cell viability, we used the Cell Counting Kit-8 (CCK-8, Selleckchem). The CCK8 solution was diluted tenfold in complete RPMI medium and then added to the patient-derived OC cells in 96-well plates. The plates were incubated for 90 minutes according to the manufacturer’s instructions. Finally, the absorbance was read at 450 nm using a Varioskan LUX microplate reader (ThermoFisher Scientific).

#### ATF-2 and P-ATF-2 Expression Analysis by Cytell Imaging System

All Patient-Derived OC Cells were seeded in 96-well plates overnight at a density of 5000 cells per well and subsequently exposed to several concentrations of CAE (20ug, 40, and 80ug/ml), then incubated for 72h. Exposed cells were then washed in PBS, fixed with 3% PFA and stained for either ATF-2 and/or p-ATF-2 (Santa Cruz) and nuclei (Hoechst). Cells were scanned and analysed using the Cytell™ imaging system (38,39).

#### Real-Time Metabolic Analysis

The oxygen consumption rate (OCR) of four patient-derived OC models (OCAS12, OCAS14, OCAS17, and OCAST16) was assessed in real-time using the Resipher system® (40). Briefly, cells were seeded in 96-well plates at 5,000 cells per well and incubated overnight. Then the baseline level of OCR was measured for an additional 24h. Subsequently, the cultured medium was replaced with fresh medium containing CAE at concentrations of 20, 40, or 80 µg/mL. OCR measurements were recorded continuously for 72 hours post-treatment. An overview of this experimental flow is shown in Figure 1.

#### Lysosomal Mass/Acidity and mitochondrial membrane potential measurements

In this study, patient-Derived OC cells (OCAS12, OCAS14, OCAS17 and OCAST16) were seeded in 96-well plates overnight at a density of 5000 cells per well and subsequently exposed to several concentrations of CAE (20, 40, or 80 µg/mL), then incubated for 72h. Following exposure to CAE cells were fixed with 3% PFA, washed in PBS and then imaged using an inverted fluorescent microscope, and changes in mitochondrial membrane potential (MMP) and lysosomal mass/pH were scanned and analysed using the Cytell™ imaging system (38,39).

#### Statistical Analysis

All the raw data from the investigated biological parameters were analysed using GraphPad Prism 8. All treatments were compared to the untreated cells for statistical significance. Statistical significance was determined using one-way ANOVA coupled with a nonparametric Tukey’s post hoc test multiple comparison test for “*” for p<0.05, “**” for p<0.01, and “***” for p<0.001. The data is presented as the mean ± standard error of the mean (n=3), and statistical significance was defined by p<0.05.

## Results

### Assessment of cancer-associated and EMT markers in Patient-Derived OC cells

Patient-Derived OC cells were characterised for a number of cancer-associated and epithelial–mesenchymal transition (EMT) markers by immunocytochemistry, and the results are shown in Figure 2.

**Figure 2:**
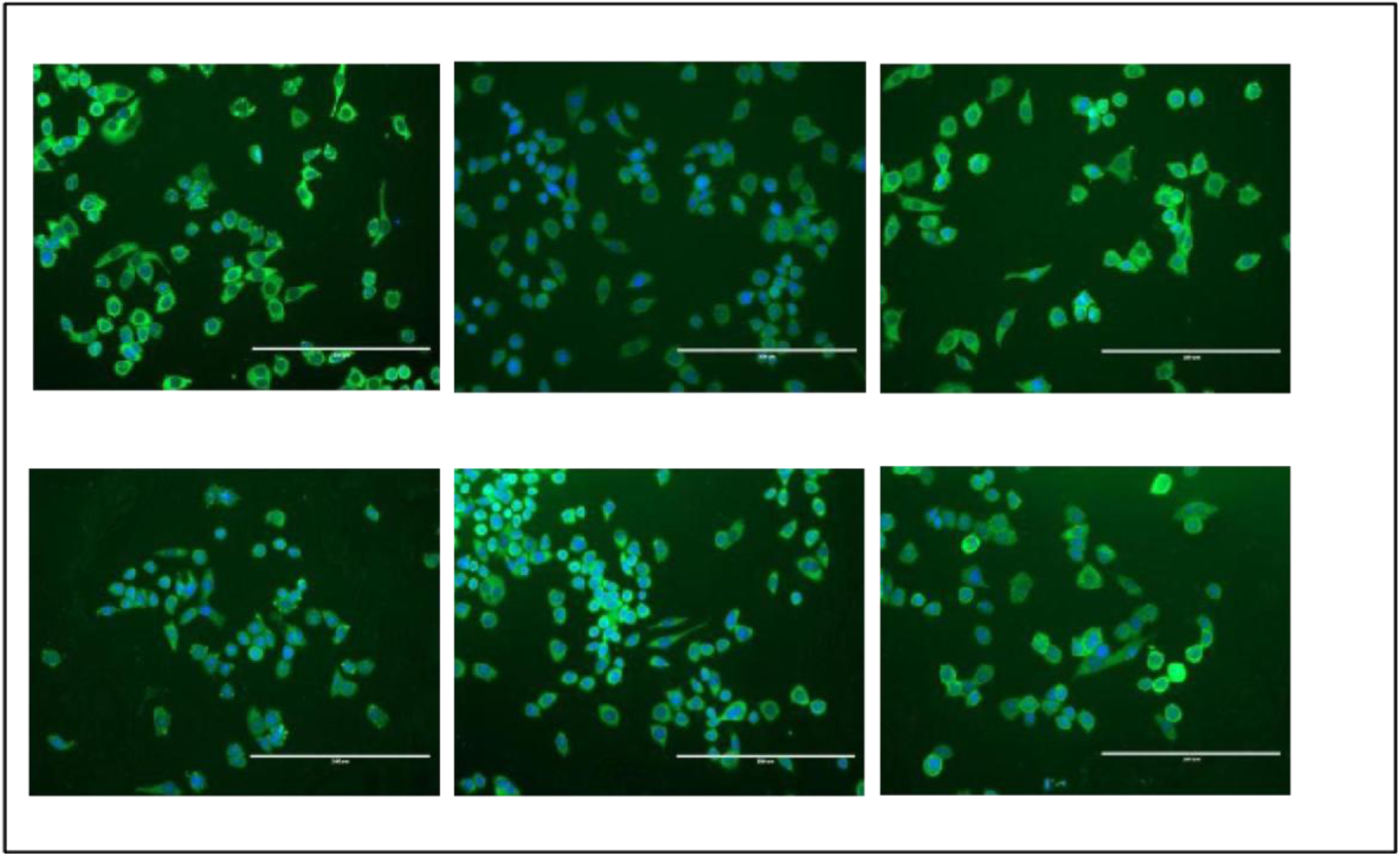
Patient-Derived OC Cells and Phenotypical Characterisation. OC cells established from the ascites samples were phenotypically profiled by cancer markers and EMT markers, including human epidermal growth factor receptor 2 (HER2/Neu), folate receptor alpha (FRα), EGFR, βcatenin, -Claudin7 and Mucin-1.

### Patient-Derived OC Models Display Distinct Sensitivity Patterns to Clove Extract

The response of OCAS12 cells to CAE showed a clear time-dependent pattern. At 24 hours, the CCK-8 assay revealed a robust dose-response relationship (Figure 3a): all concentrations (20, 40, and 80 µg/mL) significantly reduced cell viability compared with untreated controls (all p <0.001). The 40-µg dose produced a significantly greater effect than 20 µg (p <0.01) but increasing the dose from 40 µg to 80 µg offered no additional benefit, indicating an early plateau. By 72 hours (Figure 3b), this profile shifted markedly; the lowest concentration (20 µg) produced a much stronger reduction in viability, nearly triple that seen at 24 hours and the previous dose-dependency disappeared entirely. At this later time point, 20 µg performed equivalently to 40 µg and 80 µg (all p = ns), suggesting that prolonged exposure saturates the cytotoxic effect. OCAS14 cells demonstrated the opposite path. After 24 hours, all doses of CAEs significantly reduced viability relative to the control (p <0.001), but the concentrations were indistinguishable from one another, suggesting immediate maximal efficacy (Figure 3c). At 72 hours, however, OCAS14 cells developed a strong dose-response pattern (Figure 3d). Both 40 µg and 80 µg achieved significantly greater cytotoxicity than 20 µg (p <0.001), though the effect again plateaued between the top two doses (p = ns).

**Figure 3.**
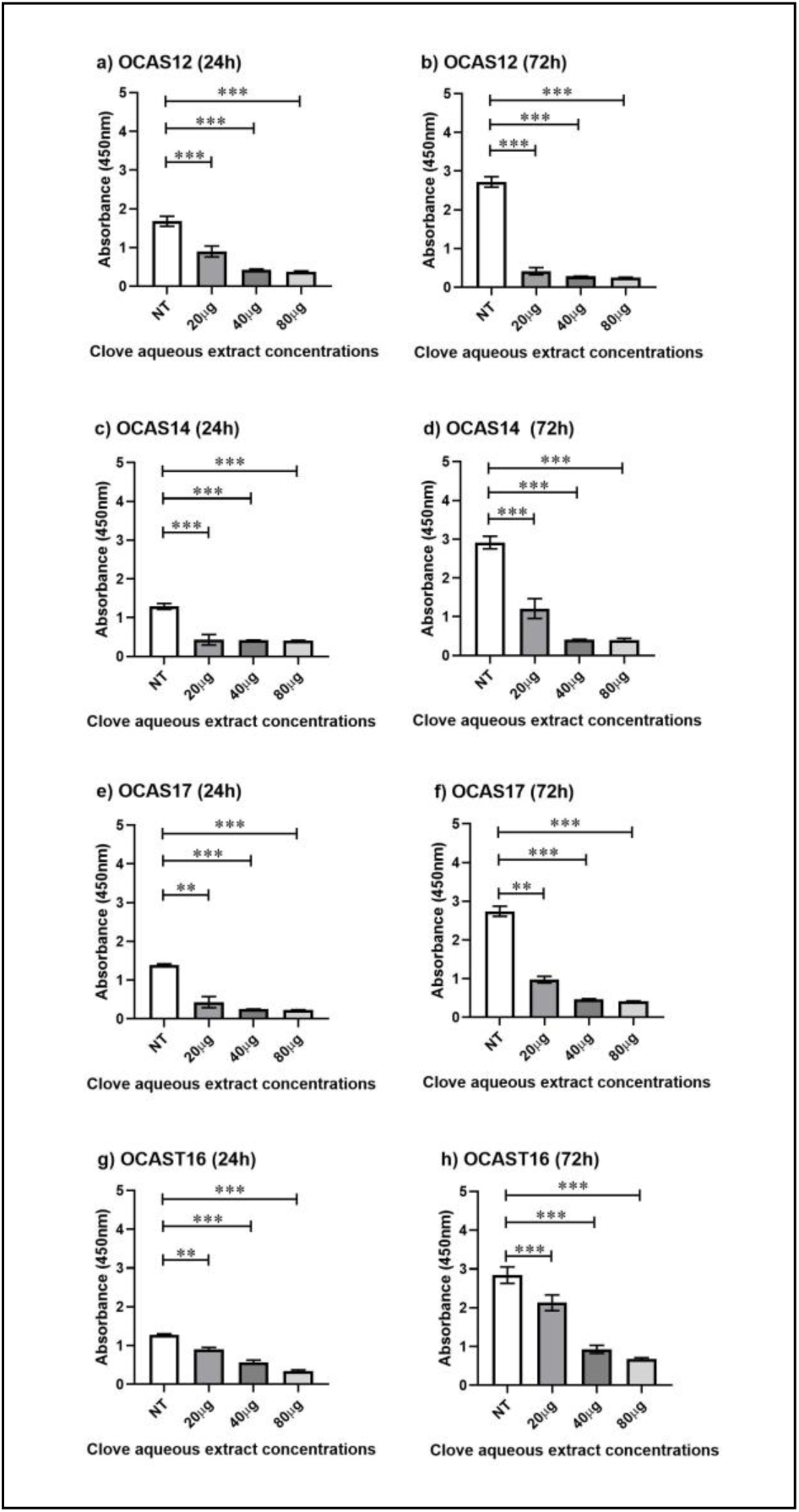
Cytotoxic Responses to CAE in Patient-Derived OC cells. Panels (a-h) show cell viability assessed by CCK-8 assay at 24- and 72-hours for four distinct patient-derived OC Cells (OCAS12, OCAS14, OCAS17, OCAST16). Cells were treated with no treatment control (NT), 20, 40, and 80 µg/mL. Data are presented as mean ± SEM of n=3 independent experiments. Statistical significance is indicated as *p < 0.05, **p < 0.01, ***p < 0.001 versus control; brackets indicate significance between doses.

OCAS17 cells exhibited an intermediate profile. At 24 hours, all doses significantly reduced viability, but only the comparison between 20 µg and 80 µg reached significance (p <0.05), indicating an early partial plateau (Figure 3e). By 72 hours, a clear dose-response emerged; both 40 µg and 80 µg were significantly more effective than 20 µg (p <0.001), although the difference between 40 µg and 80 µg remained non-significant (Figure 3f). OCAST16 cells showed a more consistent behaviour across time. At both 24 and 72 hours (Figure 3 g&h), the CAE produced a clear dose-response relationship, with each concentration significantly more cytotoxic than the last (all p <0.001). The only exception occurred between 40 µg and 80 µg at 72 hours, where the effect reached a plateau (p = ns), suggesting that 40 µg represents an optimal prolonged-exposure dose. Together, the four patient-derived models exhibited strikingly contradictory response profiles. OCAS12 lost dose-dependency over time, while OCAS14 gained it only after prolonged exposure. OCAST16 displayed consistent doseresponsiveness at both time points, whereas OCAS17 showed a gradual strengthening of dose dependence.

### CAE Differentially Regulates ATF2 Expression and Activates ATF2 Stress Signalling

CAE exerted distinct, model-specific effects on total ATF2 expression while consistently activating ATF2-mediated stress signalling across all four OC models. In OCAS12 cells, total ATF2 levels were strongly and dose-dependently suppressed at all concentrations (all p < 0.01), with clear stepwise reductions between 20 µg and 40 µg and between 20 µg and 80 µg; however, suppression plateaued between the two highest doses (Figure 4a). OCAS14 cells exhibited a similar profile, showing pronounced ATF2 suppression across all concentrations with preserved dose-dependent differences at the lower end of the range (Figure 4b). OCAS17 cells demonstrated a weaker but statistically significant reduction in ATF2 expression at all doses (Figure 4c), with significance increasing in a concentration-dependent manner (20 µg, p < 0.05; 40 µg, p < 0.01; 80 µg, p < 0.001). In contrast, OCAST16 cells uniquely displayed a paradoxical increase in total ATF2 expression at all concentrations (Figure 4d), suggesting the engagement of a distinct compensatory or adaptive regulatory mechanism that differentiates this model from the suppressive ATF2 profiles observed in OCAS12, OCAS14, and OCAS17.

**Figure 4.**
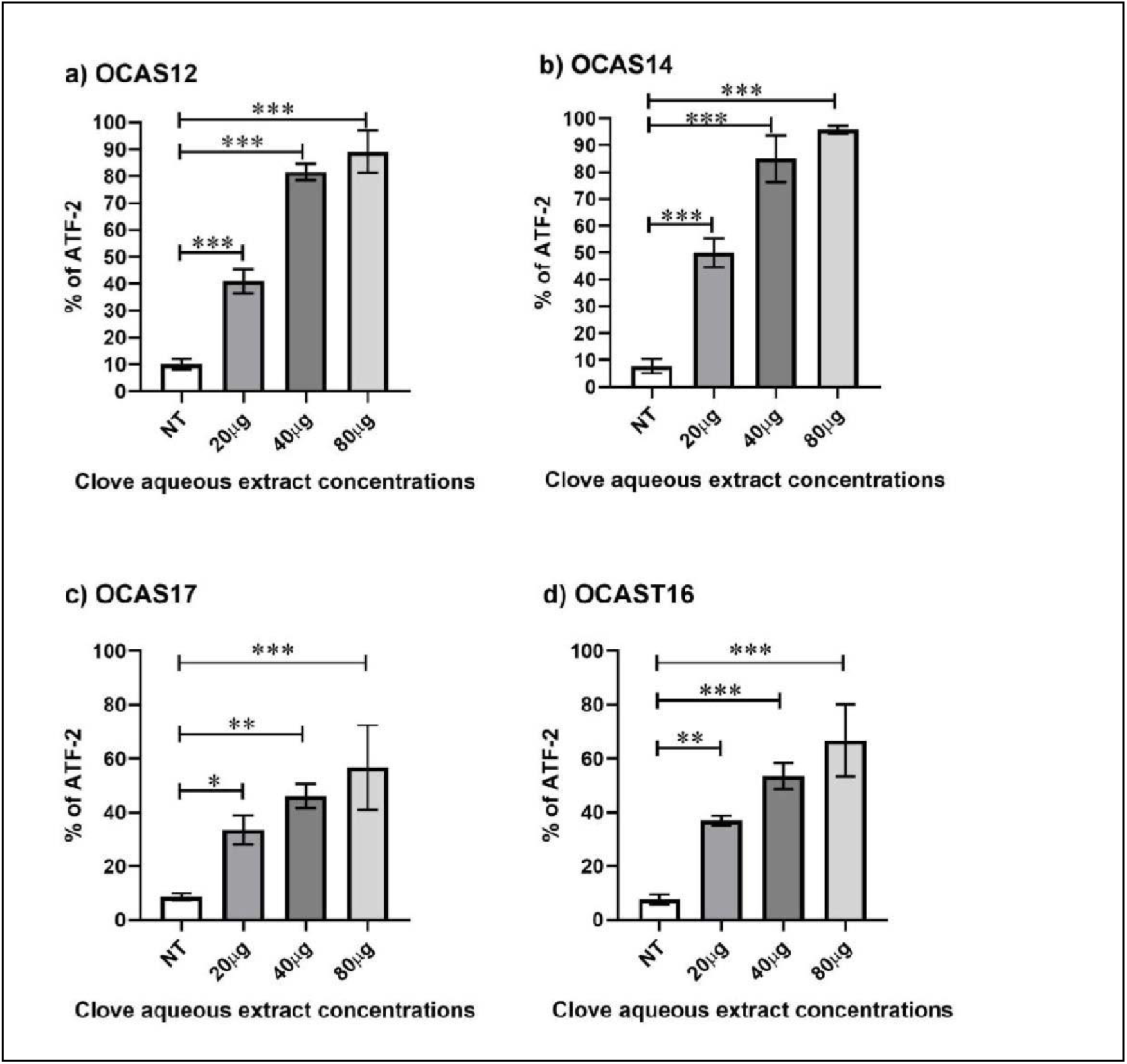
Modulation of ATF2 Expression by CAE in OC Models. Analysis of ATF2 protein or mRNA levels in four patient-derived OC cells (OCAS12, OCAS14, OCAS17, OCAST16) following 24-hour treatment with CAE (0, 20, 40, 80 µg/mL). Data are presented as mean ± SEM of n=3 independent experiments. Statistical significance was determined by one-way ANOVA with a post-hoc Tukey test. *p < 0.05, **p < 0.01, ***p < 0.001, ns = not significant.

Despite these divergent effects on total ATF2 abundance, CAE consistently increased phosphorylated ATF2 (P-ATF2) levels across all models, although the magnitude and dose sensitivity varied. OCAS12 cells showed a robust, concentration-dependent increase in PATF2 (Figure 5a) at all doses (all p < 0.001), with the response plateauing at the highest concentration. OCAS14 exhibited the most pronounced activation (Figure 5b), maintaining statistically significant discrimination even between higher doses (40 µg vs 80 µg, p < 0.01). OCAS17 cells required higher concentrations to elicit significant P-ATF2 induction; 20 µg had no effect, whereas both 40 µg and 80 µg produced significant increases (Figure 5c). In OCAST16 cells, P-ATF2 levels increased steadily across concentrations, with significant differences observed between the lowest and highest doses (Figure 5 d). Collectively, these findings demonstrate that CAE broadly engages ATF2-mediated stress signalling across patient-derived OC models, independent of its effects on total ATF2 protein levels, and highlight cell-type-specific thresholds that govern the balance between ATF2 suppression and activation.

**Figure 5.**
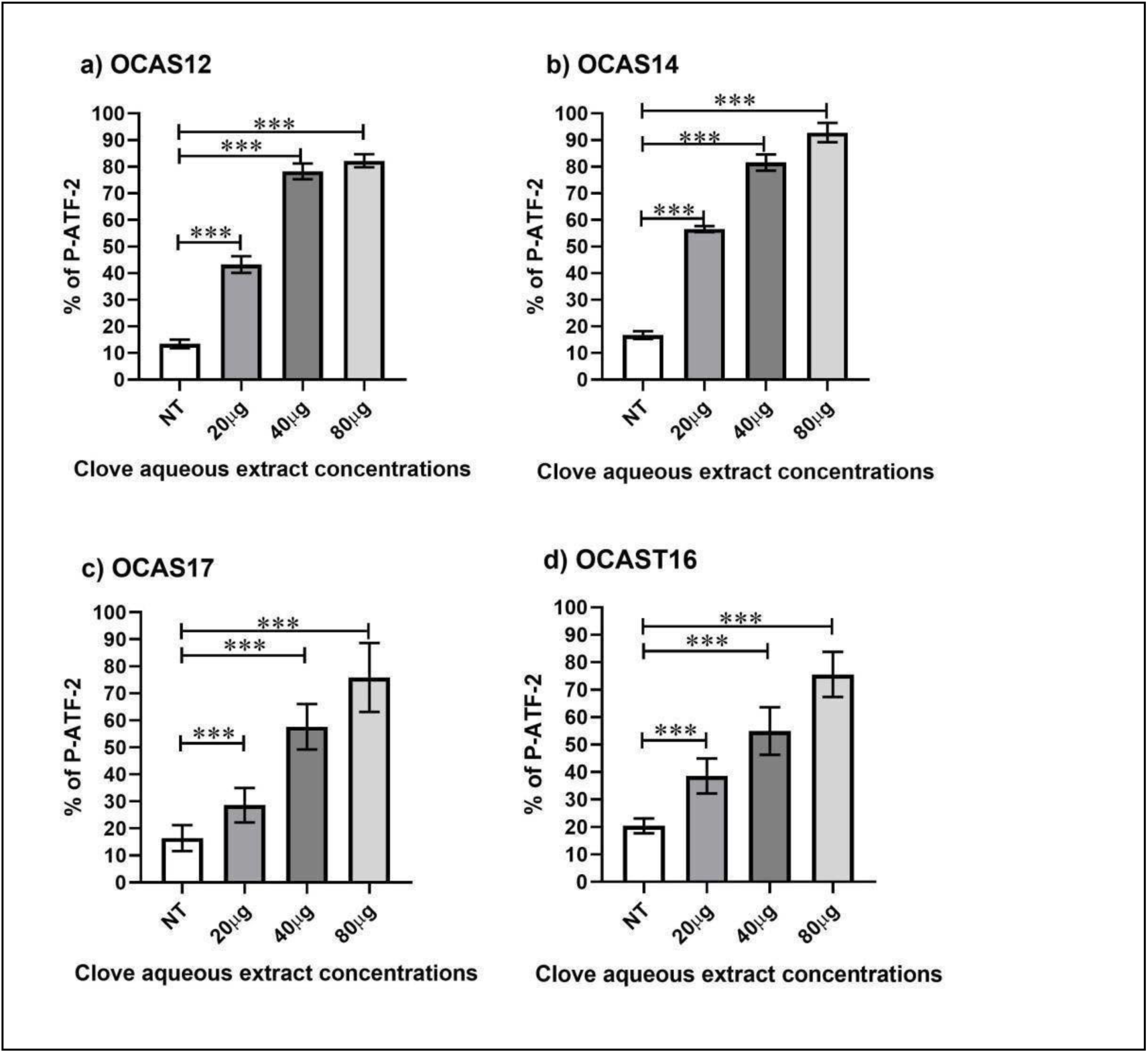
CAE activates the stress-responsive P-ATF-2: Quantification of P-ATF-2 level in **(A)** OCAS12, **(B)** OCAS14, **(C)** OCAS17, and **(D)** OCAST16 cells following treatment with CAE. Data are normalised to the non-treated (NT) control and presented as mean ± SEM of n=3 independent experiments. Statistical significance was determined by one-way ANOVA with a post-hoc Tukey test. *p < 0.05, **p < 0.01, ***p < 0.001, ns = not significant.

### CAE Disrupts Mitochondrial Metabolism in 2D and 3D Models

In two-dimensional (2D) cultures, CAE impaired mitochondrial oxidative phosphorylation across all models, as measured by OCR (Figure 6 a-h). OCAS12 cells exhibited dosedependent OCR suppression (Figure 6 b), with a 17.24% reduction at the lowest concentration (p < 0.05). OCAS14 showed a more modest response (Figure 6 d), with statistical significance achieved only at 20 µg (p < 0.05). In contrast, OCAS17 was highly sensitive, showing a 27.70% reduction in OCR at 20 µg and near-complete suppression of mitochondrial respiration at higher concentrations (Figure 6 f). OCAST16 was comparatively resistant, displaying significant effects only when compared with untreated controls (Figure 6 h). Longterm metabolic effects were further evaluated in three-dimensional patient-derived organoids (PDOs) over 8 days. CAE induced a model-dependent hypermetabolic response characterised by early metabolic stimulation followed by energetic collapse, consistent with progressive mitochondrial dysfunction (Figure 7). Ascites-derived high-grade serous ovarian cancer (HGSOC) organoids (OCAS12 and OCAS14) responded earliest (Figures 8 &9), at Days 3 and 4, respectively (all p < 0.001), whereas tumour tissue–derived OCAS16 (Figure 10) and carcinosarcoma-derived OCAS17 (Figure 11) organoids showed significant responses by Day 5 (p < 0.001 or p < 0.01). Although the timing of response initiation converged, the metabolic curves diverged substantially. OCAS12 exhibited a transient metabolic peak followed by a decline, whereas OCAS14, OCAS16, and OCAS17 displayed sustained, exponential escalation indicative of uncontrolled compensatory metabolism. Notably, OCAS16 and OCAS17 developed the most severe and persistent hypermetabolic states, consistent with heightened mitochondrial stress and reduced adaptive capacity.

**Figure 6.**
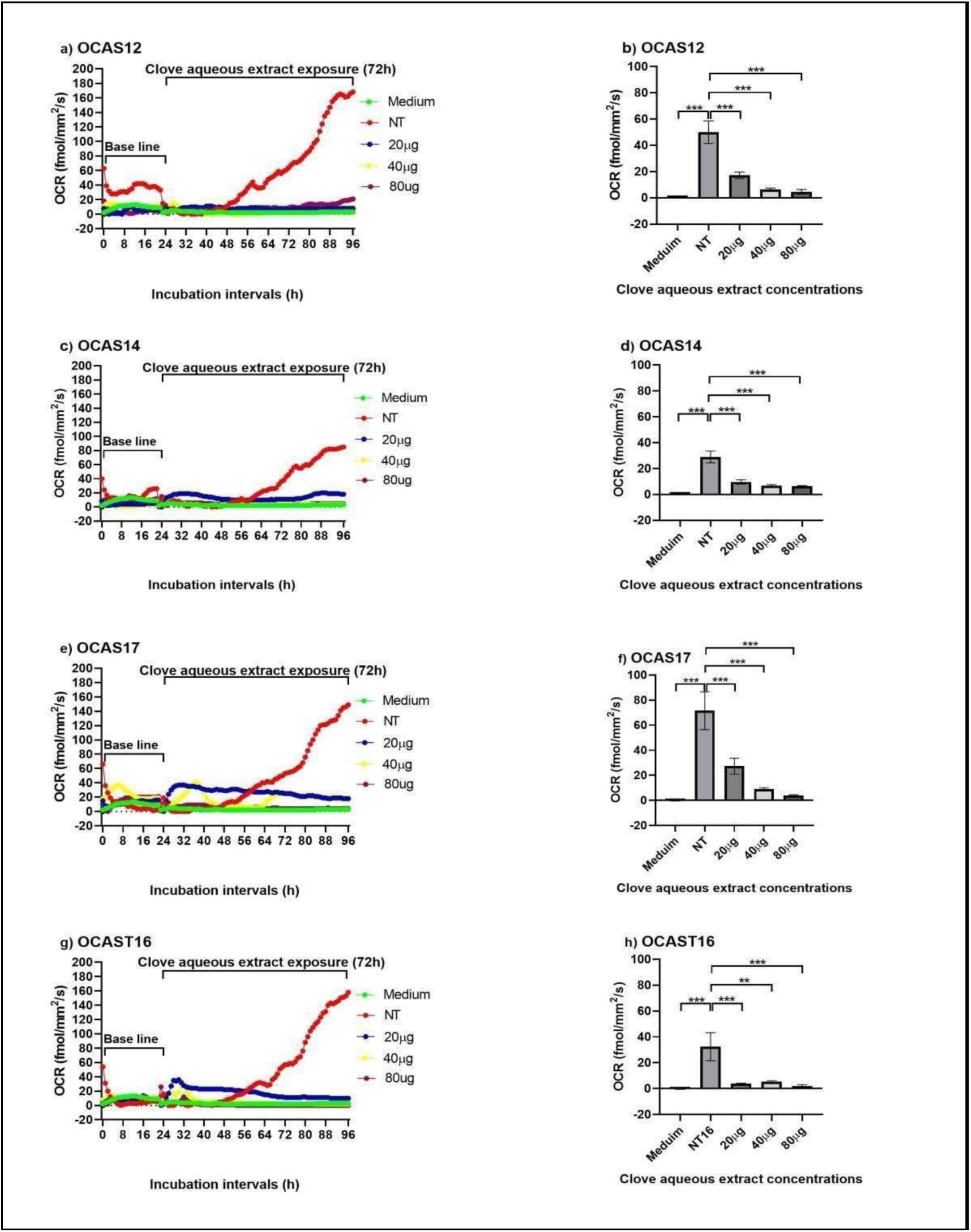
Clove aqueous extract acutely impairs mitochondrial oxidative phosphorylation in a cell line-dependent manner. Measurement of OCR, an indicator of mitochondrial oxidative phosphorylation, in **(a&b)** OCAS12, **(c&d)** OCAS14, **(e&f)** OCAS17, and **(g&h)** OCAST16 cells following 72-hour treatment with CAE. Data are normalised to the nontreated (NT) control and presented as mean ± SEM of n=3 independent experiments. Statistical significance was determined by one-way ANOVA with a post-hoc Tukey test. *p < 0.05, **p < 0.01, ***p < 0.001, ns = not significant.

**Figure 7.**
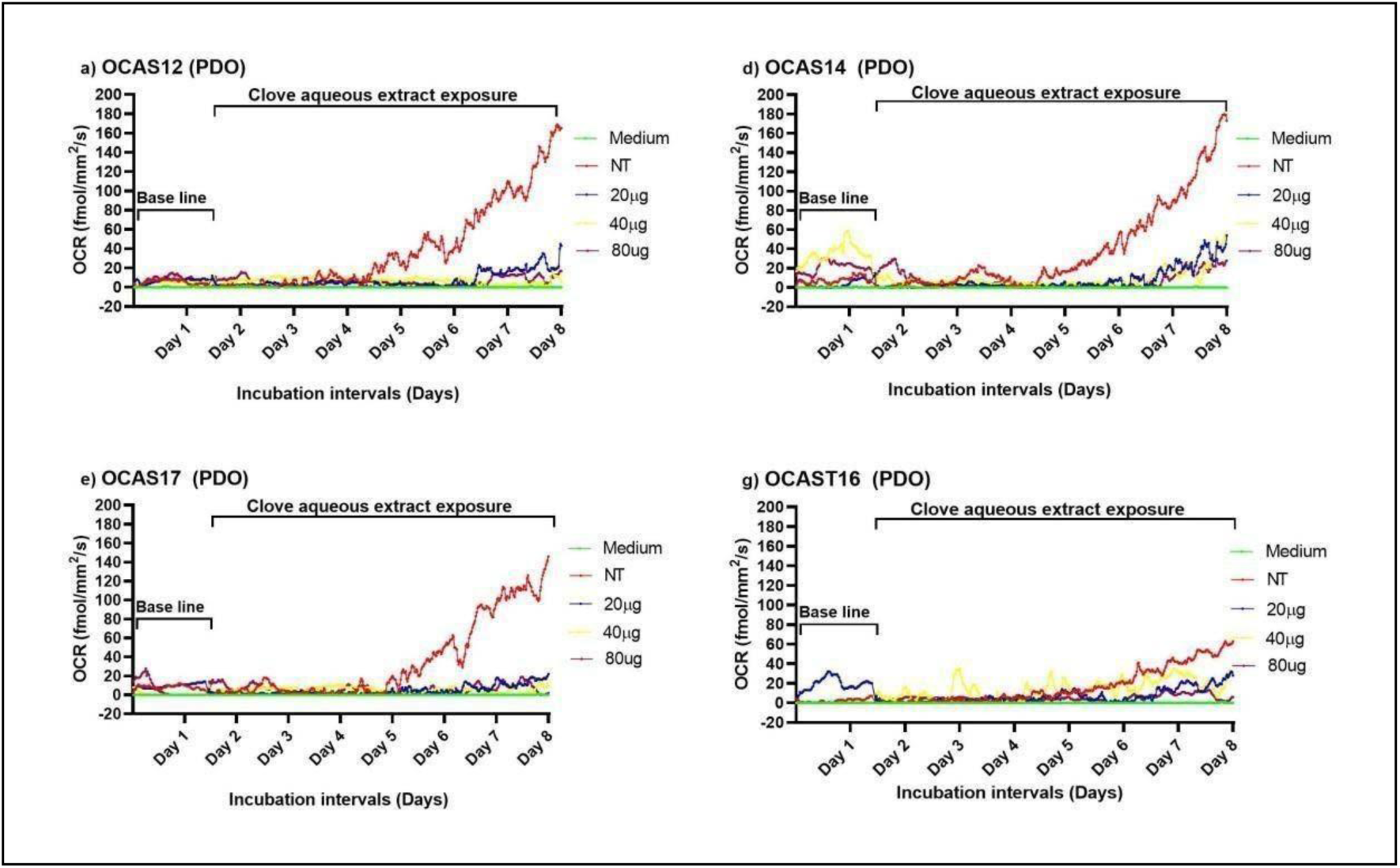
Clove aqueous extract induces a biphasic hypermetabolic response preceding energetic collapse in all PDOs (OCAS12, OCAS14, OCAS17 and OCAST16). Metabolic response curves of all OCPDOs treated with CAE over an 8-day interval, reflecting long-term treatment effects in a model that closely mimics the patient tumour microenvironment.

**Figure 8.**
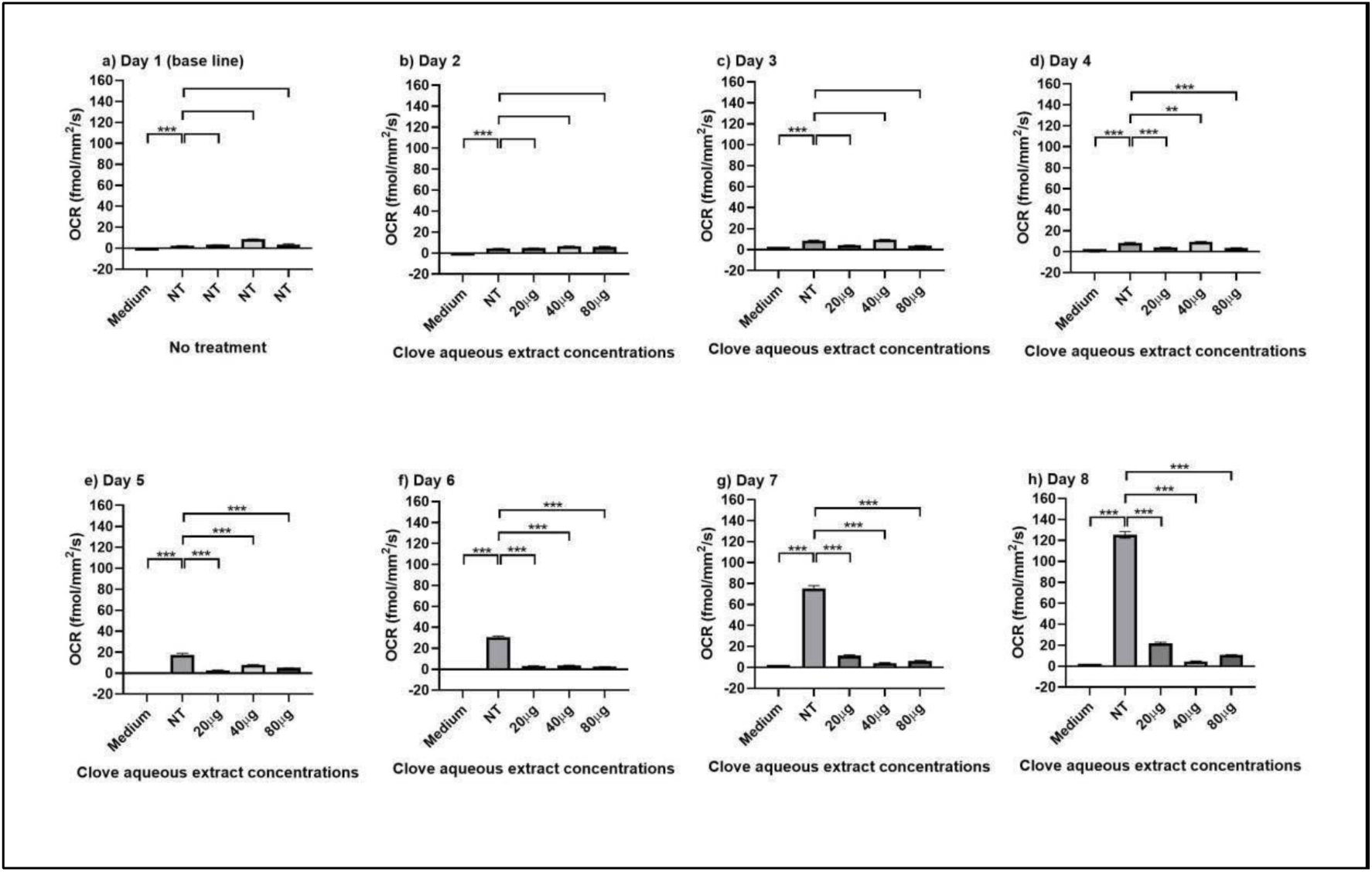
Clove aqueous extract induces a biphasic hypermetabolic response culminating in metabolic suppression in OCAS12 PDOs. Time-course analysis of metabolic activity in OCAS12 PDOs treated with CAE over 8 days. Data are normalised to the non-treated (NT) control for each time point and presented as mean ± SEM of n=4 independent experiments. Statistical significance was determined by two-way ANOVA with a post-hoc Tukey test. *p < 0.05, **p < 0.01, ***p < 0.001, ns = not significant.

**Figure 9.**
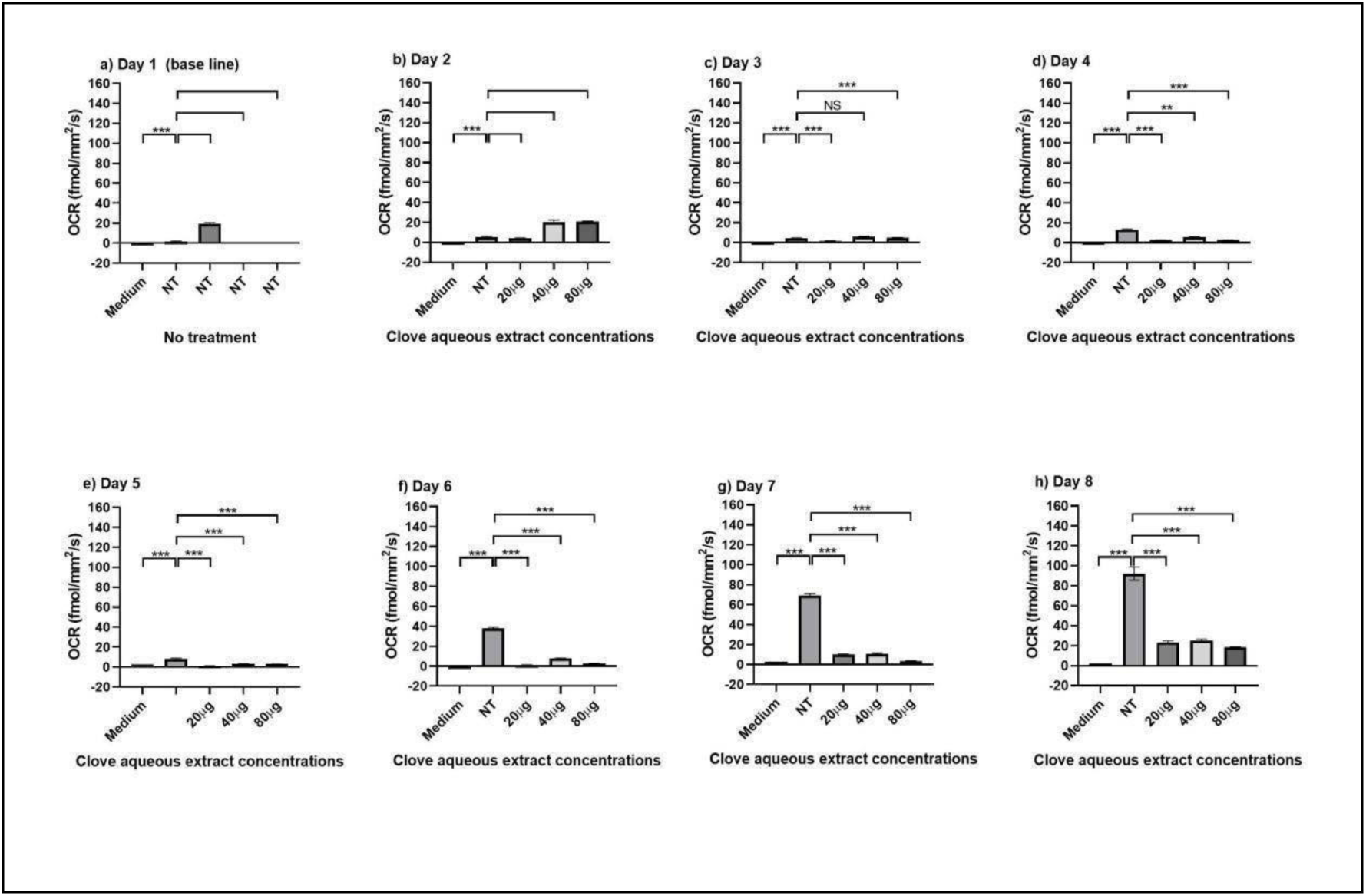
Clove aqueous extract induces a biphasic hypermetabolic response culminating in metabolic suppression in OCAS14 PDOs. Time-course analysis of metabolic activity in OCAS14 PDOs treated with CAE over 8 days. Data are normalised to the non-treated (NT) control for each time point and presented as mean ± SEM of n=4 independent experiments. Statistical significance was determined by two-way ANOVA with a post-hoc Tukey test. *p < 0.05, **p < 0.01, ***p < 0.001, ns = not significant.

**Figure 10.**
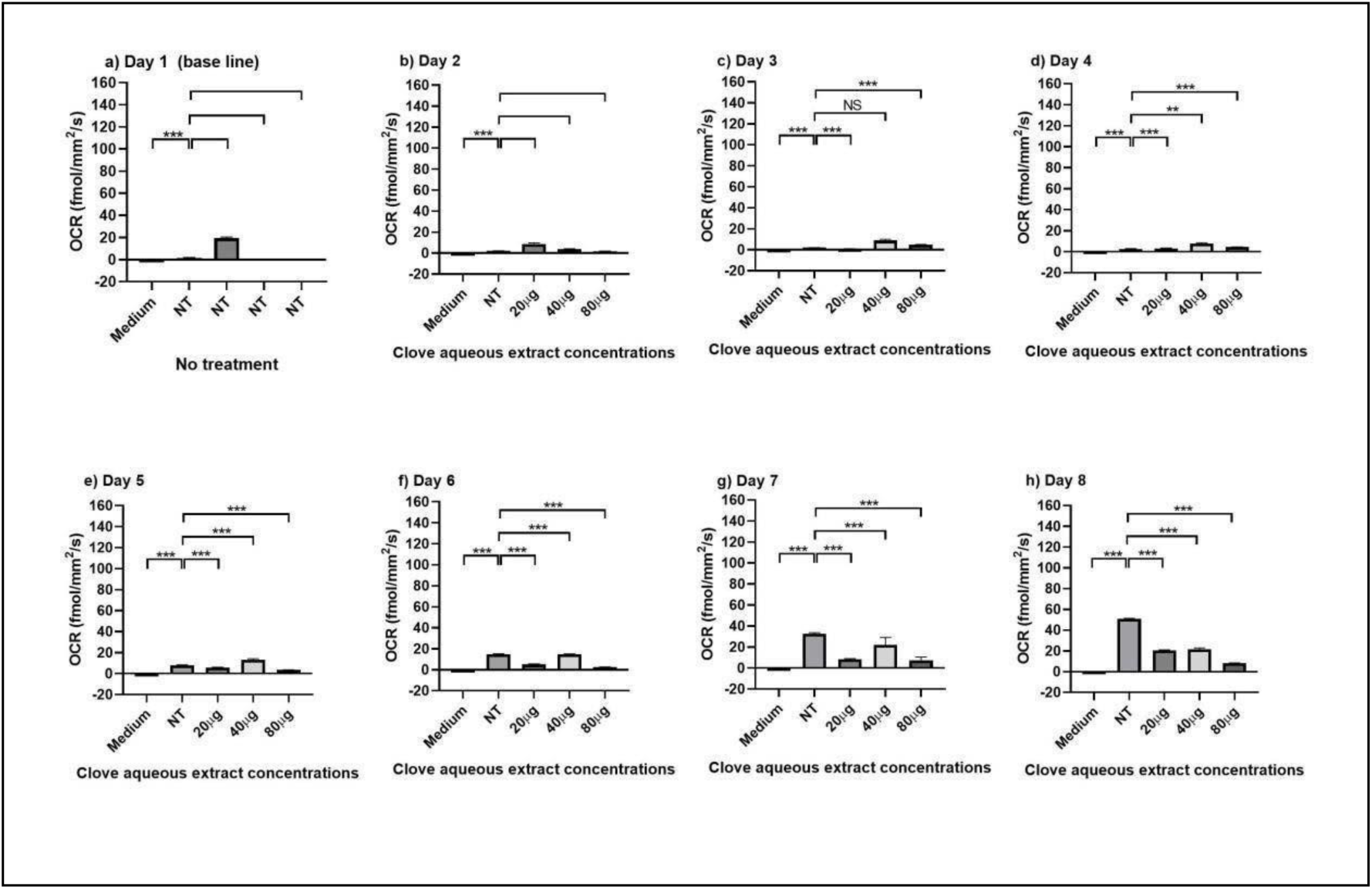
Clove aqueous extract induces a biphasic hypermetabolic response culminating in metabolic suppression in OCAS17 PDOs. Time-course analysis of metabolic activity in OCAS17 PDOs treated with CAE over 8 days. Data are normalised to the non-treated (NT) control for each time point and presented as mean ± SEM of n=4 independent experiments. Statistical significance was determined by two-way ANOVA with a post-hoc Tukey test. *p < 0.05, **p < 0.01, ***p < 0.001, ns = not significant.

**Figure 11.**
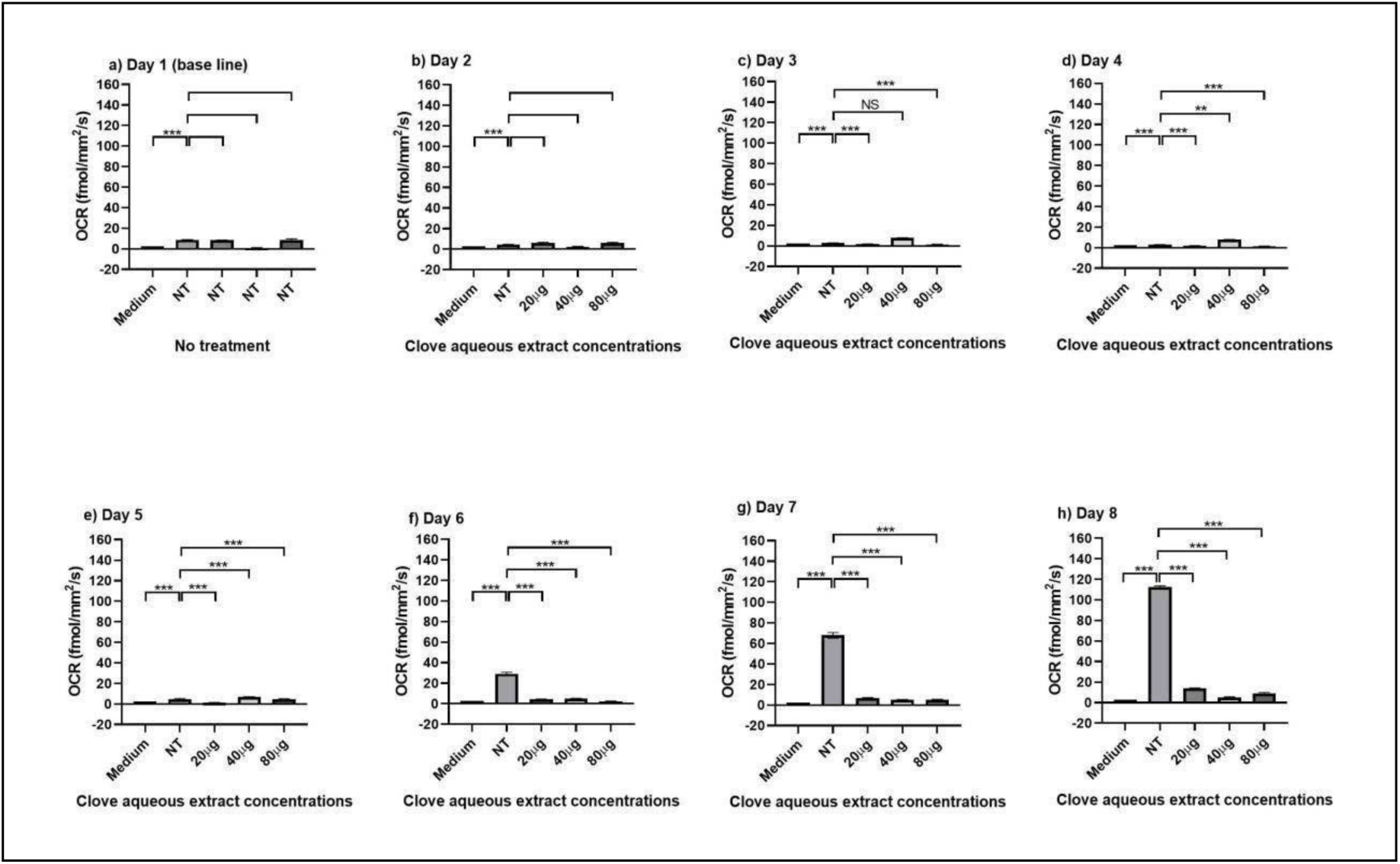
Clove aqueous extract induces a biphasic hypermetabolic response culminating in metabolic suppression in OCAST16 PDOs. Time-course analysis of metabolic activity in OCAST16 PDOs treated with CAE over 8 days. Data are normalised to the non-treated (NT) control for each time point and presented as mean ± SEM of n=4 independent experiments. Statistical significance was determined by two-way ANOVA with a post-hoc Tukey test. *p < 0.05, **p < 0.01, ***p < 0.001, ns = not significant.

### CAE-Induced Lysosomal Perturbation

It is well established that certain cytotoxic agents disrupt cellular function through several mechanisms. These include an increase in mitochondrial membrane potential (MMP), damage to the cell membrane, and impaired organelle function, specifically through the alteration of lysosomal pH or an increase in lysosome production (38, 39). To determine whether CAEinduced stress extended to the lysosomal compartment, lysosomal mass and acidity were measured across the four patient-derived OC cell lines. In OCAS12 cells, CAE induced a significant and progressive increase in lysosomal signal across all concentrations compared with nontreated controls (20 µg and 40 µg, p < 0.05; 80 µg, p < 0.01), with no significant differences between doses, indicating a robust response even at the lowest concentration (Figure 12a). OCAS14 cells exhibited a clear dose-dependent lysosomal response. While 20 µg had no effect, both 40 µg and 80 µg produced highly significant increases (p < 0.01 and p < 0.001, respectively). Significant stepwise differences were observed between 20 µg and 40 µg, 20 µg and 80 µg, and 40 µg and 80 µg, demonstrating strong concentration-dependent lysosomal perturbation (Figure 12b). Similarly to OCAS12, OCAS17 cells were highly sensitive, showing significant increases in lysosomal mass/acidity at all concentrations (20 µg and 40 µg, p < 0.01; 80 µg, p < 0.001), with effects plateauing at the lowest dose (Figure 12c). In contrast, OCAST16 cells displayed a distinct threshold response. The 20 µg dose had no effect, whereas both 40 µg and 80 µg induced sharp and significant increases (p < 0.05 and p < 0.01, respectively), confirming that a critical concentration is required to disrupt lysosomal function in this model (Figure 12d).

**Figure 12.**
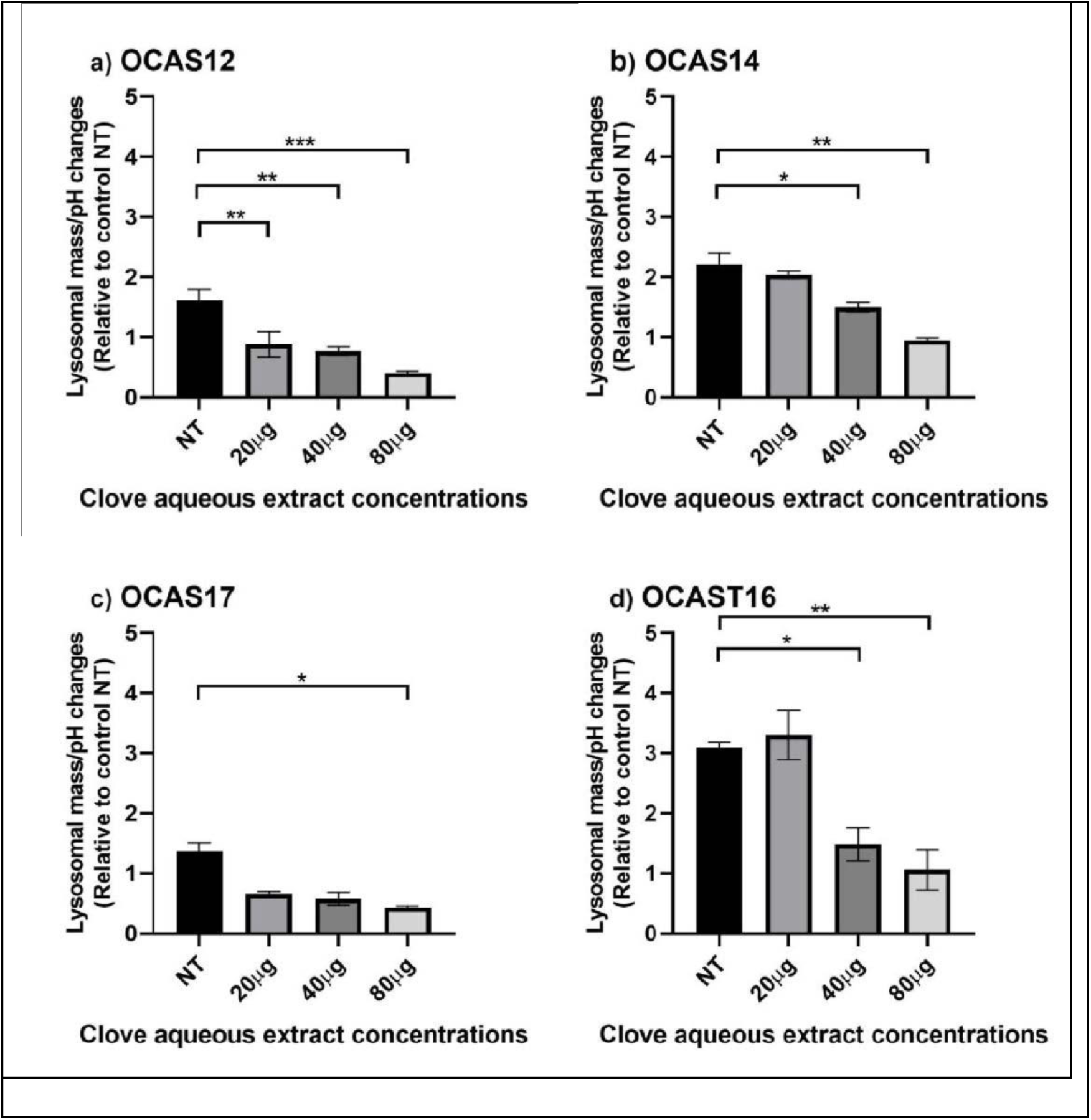
Clove aqueous extract induces lysosomal distress in a cell line-dependent manner. Quantification of lysosomal mass/pH in (A) OCAS12, (B) OCAS14, (C) OCAS17, and (D) OCAST16 cells following 24-hour treatment with clove aqueous extract. Data are normalised to the non-treated (NT) control and presented as mean ± SEM of n=3 independent experiments. Statistical significance was determined by one-way ANOVA with a post-hoc Tukey test. *p < 0.05, **p < 0.01, ***p < 0.001, ns = not significant.

### CAE Induces Mitochondrial Hyperpolarisation

Mitochondrial membrane potential (ΔΨm) was assessed to further evaluate mitochondrial stress. Rather than inducing depolarisation, CAE caused significant mitochondrial hyperpolarisation, a recognised early event in stress-induced cell death pathways. OCAS12 cells showed pronounced hyperpolarisation at all concentrations (20 µg and 40 µg, p < 0.01; 80 µg, p < 0.001), with the 80 µg dose significantly exceeding the 40 µg response (p < 0.05), indicating dose-dependent intensification (Figure 13a). OCAS14 cells exhibited dosedependent hyperpolarisation, with significant effects observed at 40 µg (p < 0.05) and 80 µg (p < 0.01), and a stronger response at 80 µg compared with 20 µg (p < 0.05) (Figure 13b). OCAS17 cells demonstrated a higher threshold for mitochondrial involvement, with significant hyperpolarisation detected only at the highest concentration (80 µg, p < 0.05), while lower doses had no effect (Figure 13c). Consistent with its behaviour in other assays, OCAST16 cells exhibited a strong threshold-dependent response, with significant hyperpolarisation induced at 40 µg (p < 0.05) and 80 µg (p < 0.01). Notably, the magnitude of this response was the greatest among all models, indicating particularly severe mitochondrial stress (Figure 13d). **Discussion** This study demonstrates that CAE exerts potent, multilayered cytotoxicity in OC through a coordinated lysosomal and mitochondrial stress program, revealing a previously unrecognised organelle-centric mechanism of action for a natural product. By leveraging patient-derived OC models, we captured clinically relevant inter-patient heterogeneity, revealing that CAE induces cell death via distinct temporal and dose-dependent pathways in different OC tumour subtypes, insights that would have been masked in conventional immortalised cell lines.

**Figure 13.**
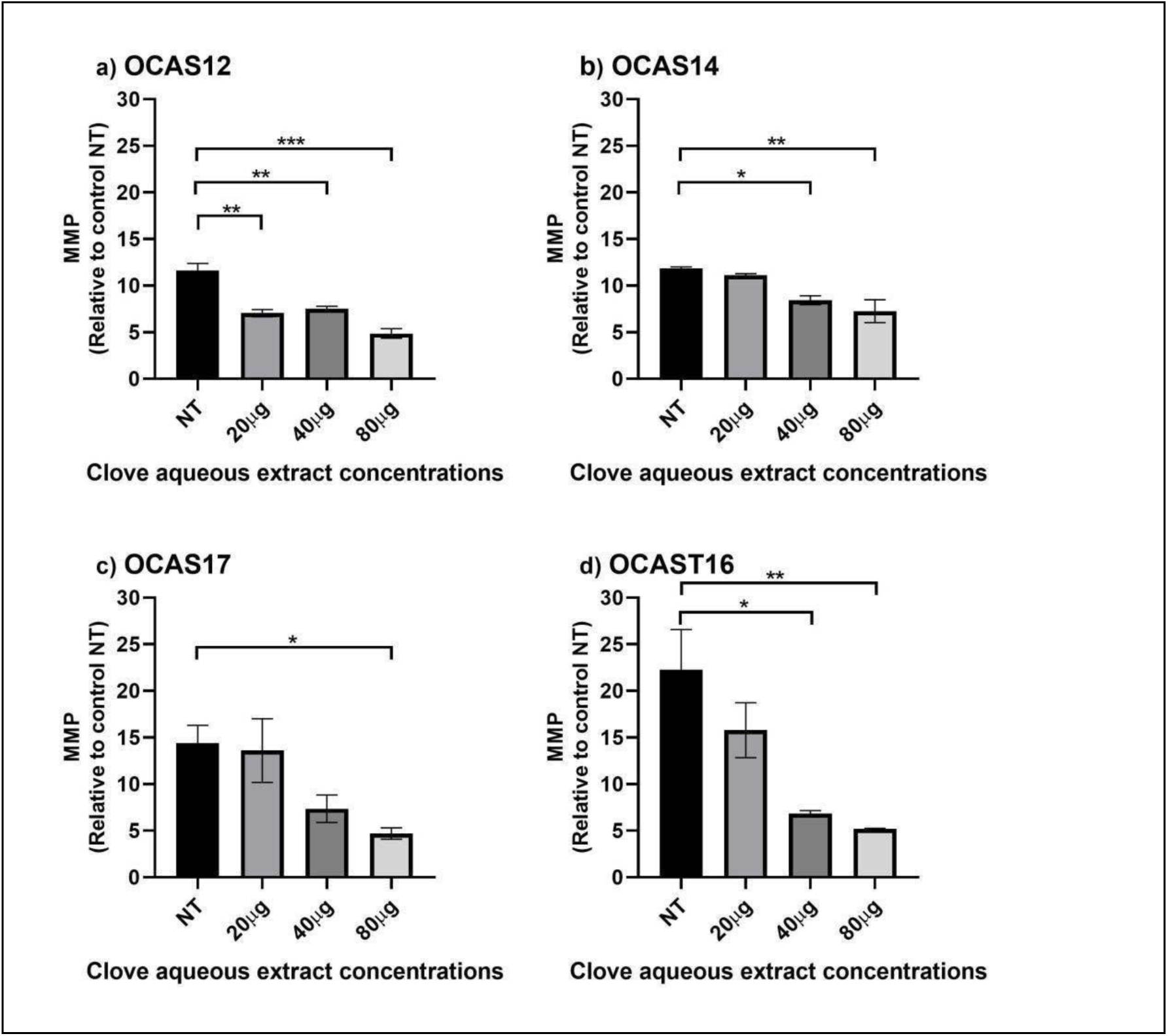
Clove aqueous extract triggers mitochondrial membrane potential hyperpolarisation. Quantification of mitochondrial membrane potential in (A) OCAS12, (B) OCAS14, (C) OCAS17, and (D) OCAST16 cells following treatment with clove aqueous extract. An increase in the metric indicates mitochondrial hyperpolarisation. Data are normalised to the non-treated (NT) control and presented as mean ± SEM of n=3 independent experiments. Statistical significance was determined by one-way ANOVA with a post-hoc Tukey test. *p < 0.05, **p < 0.01, ***p < 0.001, ns = not significant.

While natural products have long contributed to anticancer drug discovery (17–33), comparatively few studies have explored their capacity to exploit organelle-specific vulnerabilities in OC. By employing PDO models, we captured clinically relevant heterogeneity that would likely be overlooked in conventional immortalised cell lines, highlighting the translational importance of modelling patient diversity (41–45).

At the molecular level, CAE differentially modulated ATF2, a stress-responsive transcription factor implicated in DNA repair, invasion, and chemoresistance (46–48). In OCAS12, OCAS14, and OCAS17, total ATF2 expression was reduced, whereas OCAST16 showed a paradoxical increase, likely reflecting a compensatory stress response rather than sustained pathway activation. Notably, all models displayed elevated phosphorylated ATF2 (P-ATF2), albeit with varying magnitude and dose sensitivity (49–51), revealing that CAE engages a conserved ATF2-mediated stress signalling axis independent of total protein levels. These findings highlight the dynamic, context-dependent regulation of stress signalling in OC and demonstrate that consistent cytotoxic outcomes can emerge from divergent molecular responses.

CAE also profoundly disrupted mitochondrial function, reducing oxidative phosphorylation and inducing mitochondrial hyperpolarisation (ΔΨm), a recognised early apoptotic event (52–62). This metabolic collapse, reflected by dose-dependent reductions in oxygen consumption rate (OCR), highlights the bioenergetic vulnerability of OC cells. Concurrently, CAE triggered lysosomal perturbations, evidenced by increased mass and acidity in a model- and dosedependent manner (63–65). The parallel disruption of lysosomal and mitochondrial compartments illustrates organelle crosstalk as a critical determinant of cell fate, consistent with emerging literature on the role of lysosome-mitochondria interactions in apoptosis and chemoresistance (66–69).

The heterogeneity in response across PDO models underscores the importance of patientstratified therapeutic strategies. OCAS12 exhibited early and robust sensitivity, saturating at lower doses, whereas OCAS14 developed a delayed dose-response, and OCAST16 required higher concentrations to engage organelle stress. OCAS17, derived from carcinosarcoma, demonstrated relative resistance, consistent with its intrinsic chemoresistant phenotype (37,70). These observations support the potential for functional precision medicine, where dose and schedule are customised to each tumour’s vulnerabilities. Importantly, CAE’s dual organelle targeting distinguishes it mechanistically from conventional chemotherapeutics, which predominantly disrupt DNA replication or microtubule dynamics (66–69). By simultaneously compromising mitochondrial metabolism and lysosomal function pathways central to chemoresistance, CAE or its purified constituents could serve as noncrossresistant agents, either alone or as sensitisers to platinum- and taxanebased therapy (71–74). Moreover, the heterogeneity observed should not be viewed as a limitation but as clinically actionable information, enabling the identification of tumours most likely to benefit from CAEbased strategies through predictive biomarkers, such as metabolic signatures, lysosomal regulators, or ATF2 activation states.

In summary, CAE exerts potent anticancer activity in OC via a coordinated lysosomal–mitochondrial stress program, producing apoptosis through context-dependent yet mechanistically convergent pathways. The PDO platform proved essential, capturing interpatient variability, informing personalised dosing, and uncovering mechanistic complexity whereby consistent cytotoxic outcomes arise from diverse molecular responses. These findings establish a framework for the development of organelle-targeted natural products in OC and highlight the translational potential of clove-derived compounds in precision oncology.

